# A computational model of altered neuronal activity in altered gravity

**DOI:** 10.1101/2024.07.30.605832

**Authors:** Camille Gontier, Laura Kalinski, Johannes Striebel, Maximilian Sturm, Zoe Meerholz, Sarah Schunk, Yannick Lichterfeld, Christian Liemersdorf

## Abstract

Electrophysiological experiments have shown that neuronal activity changes upon exposure to altered gravity. More specifically, neurons’ firing rates increase during microgravity and decrease during centrifugal-induced hypergravity. Different biophysical explanations have been proposed for this phenomenon: however, they have not been backed by quantitative analyses nor simulations. More generally, classical computational models of neurons and networks do not account for the effect of altered gravity, which limits the possibility to perform in-silico experiments and simulations. Here, we propose computational implementations for different effects of altered gravity on cellular functions, and modify existing models to account for the effect of micro- and hyper-gravity. Firstly, in line with previous experiments, we suggest that microgravity could be modeled as an increase of the voltage-dependent channel transition rates, which is assumed to be the result of a higher membrane fluidity and can be readily implemented into the Hodgkin-Huxley model. Using in-silico simulations of single neurons, we show that this model of the influence of gravity on neuronal activity allows to reproduce the observed increased firing and burst rates. Secondly, we explore the role of mechano-gated (MG) ion channels on population activity. We show that recordings can be fitted by a network of connected excitatory neurons, whose activity is balanced by firing rate adaptation. Adding a small depolarizing current to account for the activation of MG channels also reproduces the observed increased firing and burst rates. Overall, our results fill an important gap in the literature, by providing a computational link between altered gravity and neuronal activity. Starting from historical observations of the effects of gravity on cellular functions, we derived gravity-sensitive models of neurons and networks, whose predictions could be refined using future experiments.

## 1 Introduction

During spaceflight, humans experience a variety of physiological changes due to deviations from familiar Earth conditions. Specifically, the lack of gravity is responsible for a variety of effects observed in astronauts, including a loss of muscle mass and bone density as a result of musculoskeletal de-conditioning [1–3] and modifications of the vestibular and cardiovascular systems immediately upon microgravity exposure [4]. These impairments also include structural as well as functional changes of the brain: an elevated intracranial pressure possibly leading to neuro-ophthalmic anatomical changes (spaceflight-associated neuro-ocular syndrome, or SANS) [5–7], an increased total ventricular volume [7, 8], and altered neurotransmitters functions and neurotrophic factors [9].

Given the raising interest and need of conducting experiments in microgravity conditions and the cost and complexity of accessing the International Space Station (ISS), different gravity research platforms have been developed on Earth to study the impact of gravity. Different types of 2D clinostats are mostly used to simulate microgravity in the lab [10, 11]. Real microgravity can be achieved by larger gravity research platforms, including drop towers [12], sounding rockets [13, 14], and parabolic flights with large aircraft [15–17], single-engine aircraft [18], and gliders [19]. Similarly, the effect of hypergravity can be studied using centrifuges [20, 21] or an aircraft performing high-bank turns [22].

Several of these platforms have already been employed to study how neuronal activity is impacted under altered gravity. Previous electrophysiological experiments have revealed different effects, including a higher firing rate [23–25], as well as a higher intracellular calcium concentration [11], following exposure to microgravity. Similarly, experiments using Multi-Electrode Arrays (MEA) reported an increase in the firing rates of neurons during microgravity phases and opposing effects upon exposure to hypergravity [21]. These initial changes in spontaneous action potential spiking frequency were compensated within a minute time range, hinting to an adaptation mechanism.

Different phenomena have been proposed to explain the increased firing rate during microgravity and the decreased firing rate during hypergravity. First, a possible explanation is that microgravity increases the membrane fluidity of neuronal cells, which modifies properties of the ion channels. More specifically, a previously proposed model [24] hypothesizes that an increase in membrane fluidity modifies the open-state probabilities of ion channels, hence leading to a depolarization of the cell and to an increase in the spontaneous firing rate. Although intuitive, this model is purely qualitative and has not been backed by analytical computations nor simulations. In computational models, a neuron is often described by a set of parameters, e.g., their membrane time constant *τ*_*m*_ or their ion channel conductance *g*. A proper description of how these parameters might be impacted by altered gravity conditions is currently missing.

A second explanation to the increased firing and burst rates in microgravity is based on the role of mechanosensitive ion channels, which operate as gravireceptors in eukaryotic cells [26]. They have been shown to be widely present in the nervous system [27], including hippocampal [28] and pyramidal neurons [29]. These mechano-gated (MG) ion channels have been observed to generate inward currents and to elicit spiking activity in neuronal cells [29]. The modified neuronal activity during periods of altered gravity has thus been attributed to the activity of MG ion channels, whose stretching during acceleration and deceleration phases might impact neuronal activity [21]. But the final effect of an inward current on firing rates is going to strongly depend on the properties of the studied neuron population. Specifically, these MG channels would have different effects depending on whether the observed neurons are isolated or show signs of synchronized activity; whether their recurrent connections are mostly excitatory or inhibitory; or whether they are impacted by non-linear dynamical effects, such as short-term depression or firing rate adaptation. To date, these features have not been characterized in the neuron cell types used in altered gravity experiments.

First, in section 2.1, we propose that the effect of altered gravity on neuronal dynamics could be represented as an acceleration of the rate at which voltage-dependent ion channels switch from open to close configurations, i.e., as a decrease of the time constants *τ*_*n*_(*V*), *τ*_*m*_(*V*), and *τ*_*h*_(*V*). This is in line with a set of previous observations reporting a modification of the fluidity of the cell membrane in altered gravity, and a direct link between membrane lipids and ion channels. We verify that decreasing these time constants during in-silico simulations allows to reproduce the increased firing and burst rates observed during altered gravity experiments (section 2.2).

Second, in section 2.3, we propose a computational model to explain the activity of networks of neurons in microgravity. Human stem cell-derived neuronal cells which were already used in microgravity experiments have two remarkable features: they are mostly excitatory, and show slow oscillations of coordinated activity [30]. We show that these features can be accounted for by a network of connected excitatory leaky integrate-and-fire (LIF) neurons which recurrent connections are balanced by firing rate adaptation. We verify that injecting an external depolarizing current into the network (to model the activation of MG channels) also allows to reproduce the increased firing and burst rates observed during altered gravity experiments (section 2.4).

## 2 Results

### 2.1 A proposed model of the effect of gravity on ion channels’ gating rates

A classically used model of the time evolution of the membrane potential of a neuron, and especially of how action potentials are generated, is called the Hodgkin-Huxley model [31, 32]. It consists of a set of differential equations, describing the evolution of the cell voltage as a function of its ion channels conductances; the evolution of these conductances as a function of the open probabilities of the channels; and the evolution of these probabilities as a function of their voltage-dependent gating rates.

The standard equation describing the time evolution of the voltage *V* of a single-compartment model of a neuron (see [32] for a detailed derivation) is

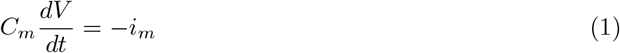

where *C*_*m*_ is the specific membrane capacitance (in F *·* m^−2^) and *i*_*m*_ is the total membrane current. The latter can be expressed as the sum of a delayed-rectified potassium current, a transient sodium current, and a leakage current:

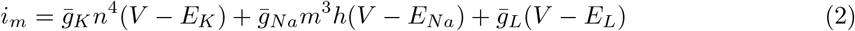

where *E*_*L*_, *E*_*Na*_, and *E*_*K*_ are the reversal potentials, and 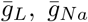, 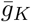, and 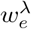 are the maximal surface conductances (in S *·* m^−2^). *n* corresponds to the probability that a subunit in a potassium channel is activated and hence contributes to the opening of the *K*^+^ channel. Similarly, *m* and *h* express respectively the activation and inactivation probabilities of the *Na*^+^ conductances. The time evolution of these gating variables can also be described by a set of differential equations:

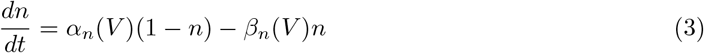

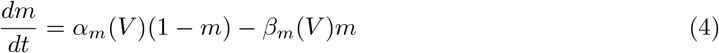

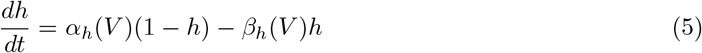

where *α*_*x*_(*V*) and *β*_*x*_(*V*) are the voltage-dependent opening and closing rates of the subunit gates, which are usually obtained by fitting on experimental observations (see [32] for a detailed discussion). Eq. 3 can also be written as

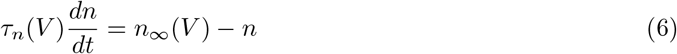

With

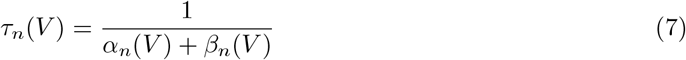

and

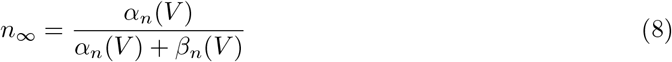

(similar equalities can be derived for the other gating variables *m* and *h*). The time constants *τ*_*n*_(*V*) (Eq. 6), *τ*_*m*_(*V*), and *τ*_*h*_(*V*) thus govern the dynamics of channel subunits, i.e., the rate at which they open and close.

Altered levels of gravity have been observed to act on the activity of single ion channels [33–35], but more importantly to act on the fluidity of the cell membrane itself. Studies using random positioning machines (RPM) reported a dramatic increase in the fluidity of cell membrane models under simulated microgravity conditions [36]. The increased fluidity under microgravity (and reciprocally the reduced fluidity during hypergravity) have been further observed in different membranes [37] and different altered gravity modalities [23, 25].

In a previously proposed model [23, 24], a link between gravity-induced changes in fluidity and firing rates was suggested: it hypothesizes that an increase in membrane fluidity modifies the properties of ion channels, hence leading to a depolarization of the cell and to an increase in the spontaneous firing rate. Although heuristic, this model is purely qualitative and has not been backed by quantitative analyses nor simulations.

However, several independent studies reported that ion channel functions are regulated by the cell membrane’s lipid microdomains [38], and that lipid binding sites exist on ion channels, yielding fluidity-dependent interactions between them [39]. More specifically, arachidonic acid has been shown to increase the rate of voltage-gated calcium channels activation kinetics [40], while lipid gating influences the activity of potassium channels [41]. Additionally, a modulation of acetylcholine receptors by membrane lipids was highlighted^1^ [43], as well as a link between medium temperature, membrane fluidity, and channels closing rate [44].

We propose to represent the effect of microgravity on computational models of neurons by a unitless multiplicative constant *λ* which is applied to the opening and closing rates *α*_*x*_(*V*) and *β*_*x*_(*V*), and hence to the time constants *τ*_*n*_(*V*), *τ*_*m*_(*V*), and *τ*_*h*_(*V*) (Fig. 5). The classical gating equations (Eqs. 3, 4, and 5) thus become

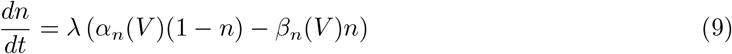

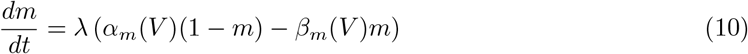

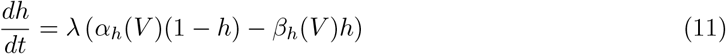

with *λ* = 1 in 1*g, λ <* 1 in hypergravity, and *λ >* 1 in microgravity. Fig. 5 shows that, for a given voltage, increasing the value of *λ* decreases the gating time constants (left panel) without modifying the gating steady-state levels (right panel). Our approach allows to reconcile two independent observations: the fact that microgravity impacts the cell membrane fluidity [23, 25, 36, 37]; and the fact that membrane lipids impact the channels functions [38–44].

### 2.2 In-silico simulations of single cell activity under altered gravity

Next, we sought to assess the effect of *λ* on simulated neuronal activity, and especially to verify that a higher value of *λ* (to represent the effect of microgravity) would reproduce the different features observed during experiments, namely a higher firing rate and burst rate - and that a value of *λ* set below 1 would reproduce the reduced activity observed following the onset of hypergravity in centrifuge experiments. Simulations were performed using the Brian neuron simulator [45] and the numerical values described in Methods.

Fig. 1A shows the firing rate of a neuron simulated by the Hodgkin-Huxley model as a function of the input current *I* added to Eq. 2 and of the value of *λ* in Eqs. 9, 10, and 11. For a given input current, the firing rate increases with *λ*, which is in line with experimental observations. We verified that this result is robust to the values used in the simulation by randomly varying the values of the neuron’s parameters and checking that a higher value for *λ* robustly leads to a higher firing rate (Fig. 6). Fig. 1B shows the time evolution of the voltage of the same neuron for an input current *I* = 0.35nA. A shorter time constant for the gating variables leads to a faster voltage increase, an earlier time to first spike, and a shorter inter-spike interval.

**Figure 1:**
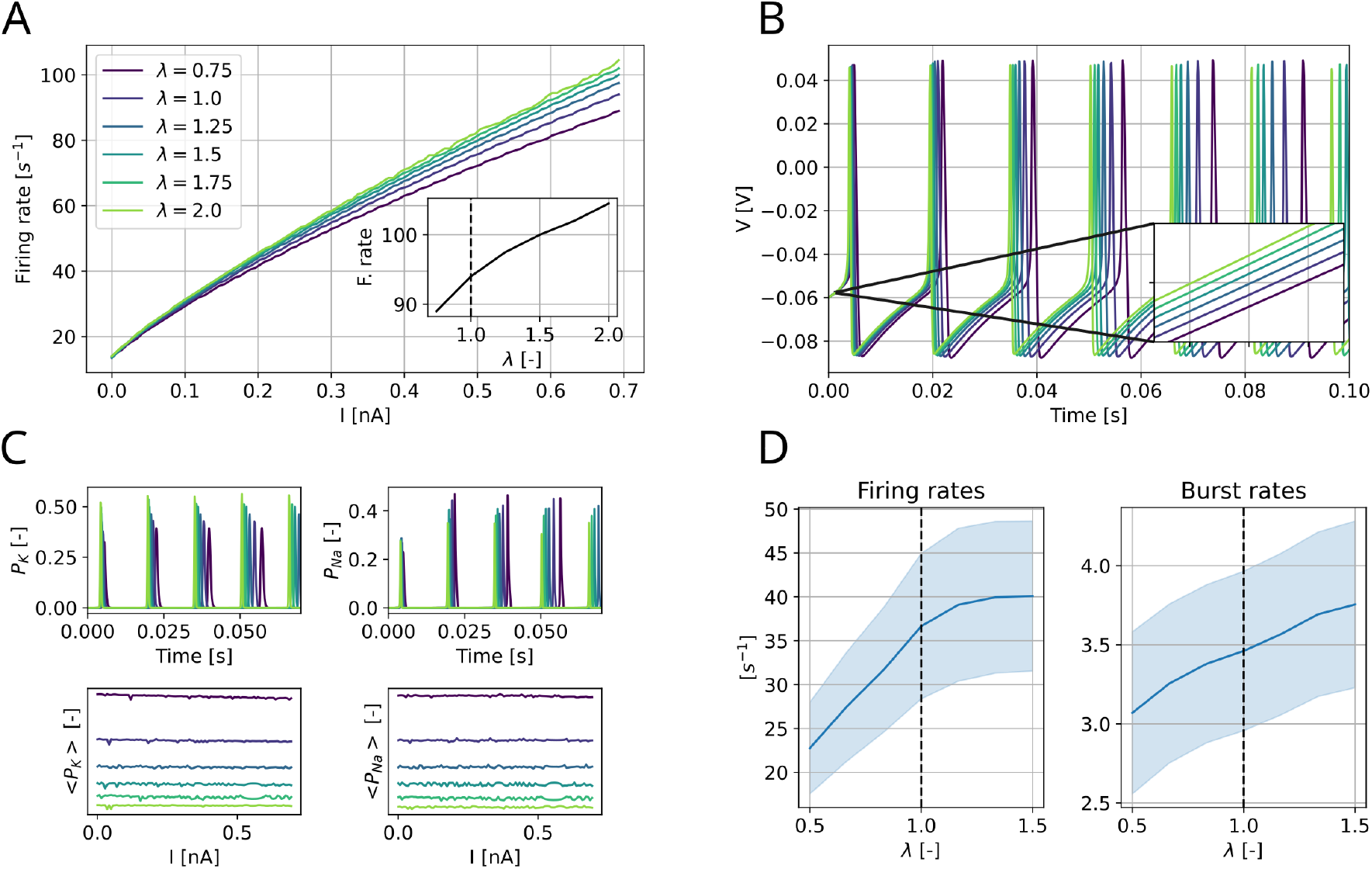
A: Frequency-current curve of a simulated Hodgkin-Huxley neuron (whose parameters are described in Table 1) for different values of *λ*. Inset: firing rate as a function of *λ* for *I* = 0.7nA. B: Time evolution of the membrane voltage for the same cell as in A. Inset: zoom on the first ms of the simulation, showing the earlier depolarization induced by a higher value for *λ*. C: Evolution of potassium *P*_*K*_ (left column) and sodium *P*_*Na*_ (right column) channel opening probabilities for the same cell as in A. Upper row: time evolution of these probabilities for *I* = 0.35nA. Lower row: average (over *T* = 2s) value of these probabilities (normalized by the number of spikes during the simulation). D: Mean firing and burst rates (averaged across all neurons) in a network of connected Hodgkin-Huxley neurons as a function of *λ*. Solid line: mean value over 30 repetitions. Shaded area: s.e.m.

Fig. 1C shows the effect of *λ* on the open probabilities of the potassium and sodium channels, i.e., *P*_*K*_ = *n*^4^ (left column) and *P*_*Na*_ = *m*^3^*h* (right column). The upper row shows their time evolution in the same condition as in Fig. 1B, while the bottom row shows the average value of *P*_*K*_ and *P*_*Na*_ during 2 seconds of simulation (normalized by the number of action potentials). The end results in the lower row is independent of *I*, since the firing rate is nearly linear with *I* for sufficiently large values (see Fig. 1A). These results show that, under our modeling assumptions, an increase in *λ* does not increase the opening probabilities of the ion channels, but rather accelerates the time at which the *Na*^+^ conductances activate. Previous studies had reported seemingly contradictory results on the effect of altered gravity on ion channels opening probabilities [33–35]. Additionally, previous qualitative models of the increased firing rates observed in microgravity were based on the assumption that microgravity exposure would reduce the ion channels opening probabilities, ultimately yielding an increased firing rate [24]. However, ion channels are dynamical systems, for which the opening probability cannot be reduced to a single time-invariant value. Furthermore, they are not isolated, and their dynamics are voltage-dependent and correlated to the activity of other channels in the cell. Here, we suggest a more nuanced approach to the effect of gravity on ion channels, in which altered gravity does not simply modify the baseline opening probabilities of channels but rather acts on their dynamics.

**Table 1:**
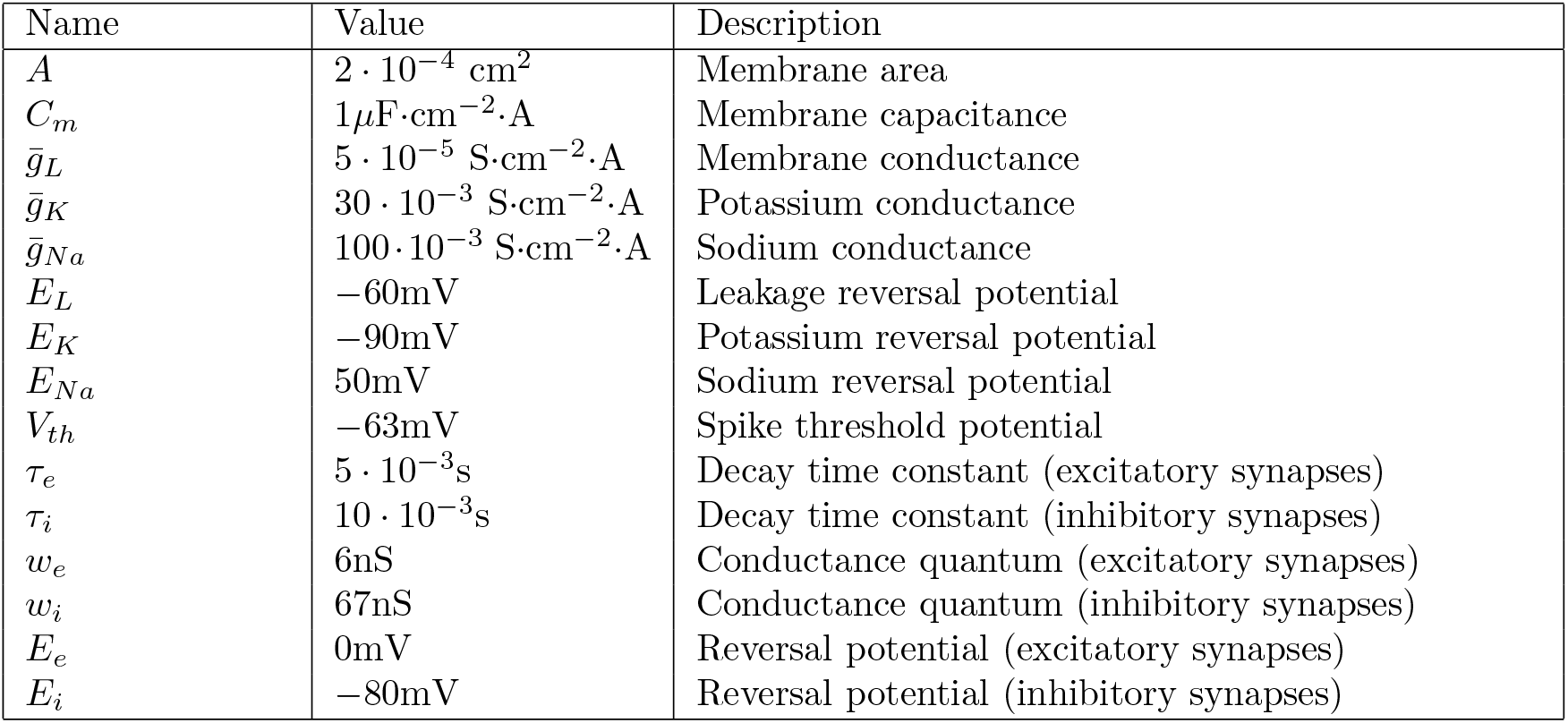
Parameter values used in Fig. 1 (similar values as in [67]). The detailed equations for the Hodgkin-Huxley model and the COBA synapses used in Fig. 1 are also described in [67].

Finally, we also verified that varying the value of the parameter *λ* allows to reproduce experimental observations not only in isolated neurons, but also in more realistic networks of recurrently connected neurons. To do so, we simulated the activity of a population of *N* = 4000 connected Hodgkin-Huxley neurons (whose parameters are described in Table 1), and verified that increasing the value of *λ* leads to an increase of both the population firing rate (Fig. 1D, left), and burst rate (Fig. 1D, right). In line with previous studies [21], we define a burst as a group of consecutive spikes separated by less than 20 ms. These results are akin to what has been previously observed during altered gravity experiments [21].

### 2.3 Observed synchronized bursts can be explained by spike frequency adaptation

In addition to the role of cell membrane fluidity, another possible explanation for the altered neuronal activity involves MG ion channels, whose activation in altered gravity phases would depolarize cells [21]. However, although this explanation makes sense for single isolated neurons, the effect of an inward current on recurrently connected neurons would depend on the properties of the network. More specifically, it would depend on its excitation-to-inhibition ratio, and on the presence of non-linear dynamics, such as short-term plasticity or firing rate adaptation. Such features have not been characterized, and let alone modelled, in the cell lines classically used in altered gravity experiments.

Such cell lines usually include human stem cell-derived neuronal cells (i.e. iNGN cells) (see [30] and [21] for a detailed discussion), which have two main remarkable features. Firstly, their population activity shows synchronized bursts and action potentials (Fig. 2A, and Fig. 3C in [30]) at a low frequency (i.e. *<* 1Hz), which is a telltale sign of recurrent connections among neurons. Secondly, the vast majority of these connections are supposed to be excitatory: synchronized activity in iNGN neuronal cell lines is abolished by glutamate receptor blockers, and although inhibitory GABAergic synapses appear in the last stages of cells development, inhibitory neurons only make up approximately 2.3% of the whole population after 30 days of cell growth [46]. This is significantly below the level of inhibition which is classically assumed to be necessary for stable self-sustained population activity [47].

**Figure 2:**
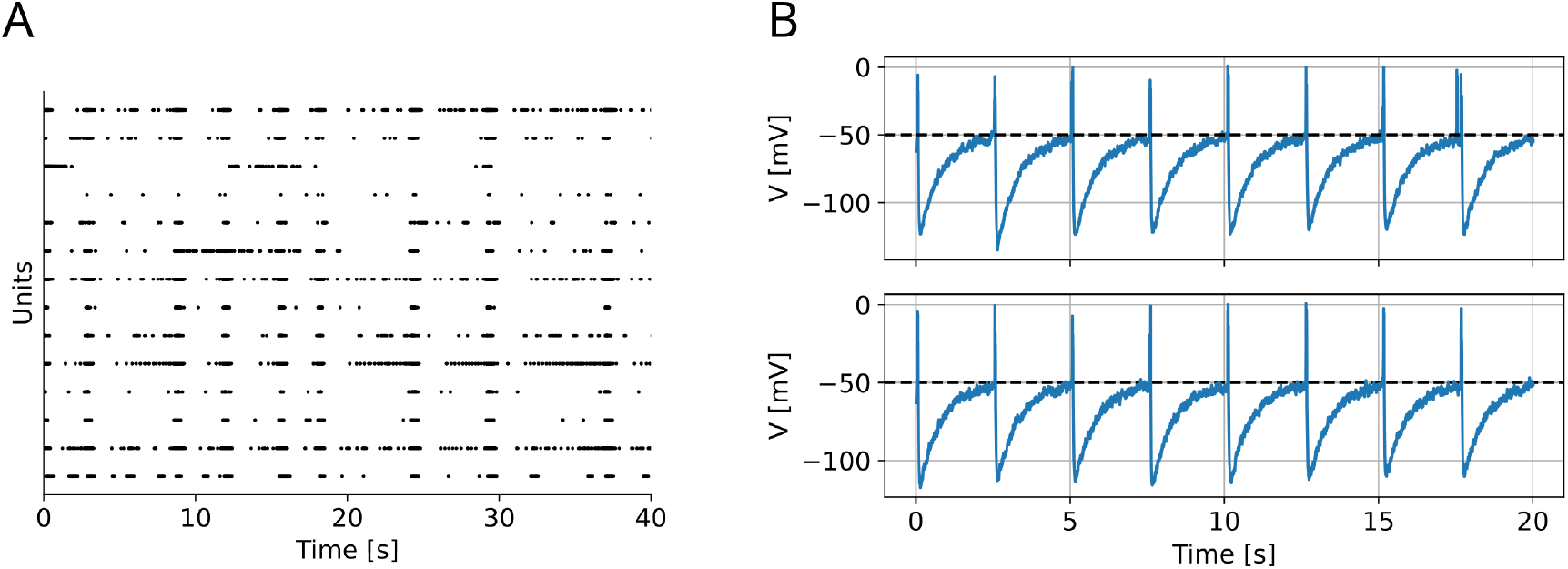
A: Raster plot of detected action potentials on 14 units observed from a human stem cell-derived neuronal network (same setting as in [21]). Spike sorting was performed using SpikeInterface [54] and the MountainSort 5 algorithm [55]. B: Membrane voltage of 2 example cells from a simulated network of connected LIF neurons (without inhibition but with firing rate adaptation). Dashed dark line: action potential threshold. Numerical values used for the simulations are described in Table 2. Bursts happen regularly approx. every 5s.

**Figure 3:**
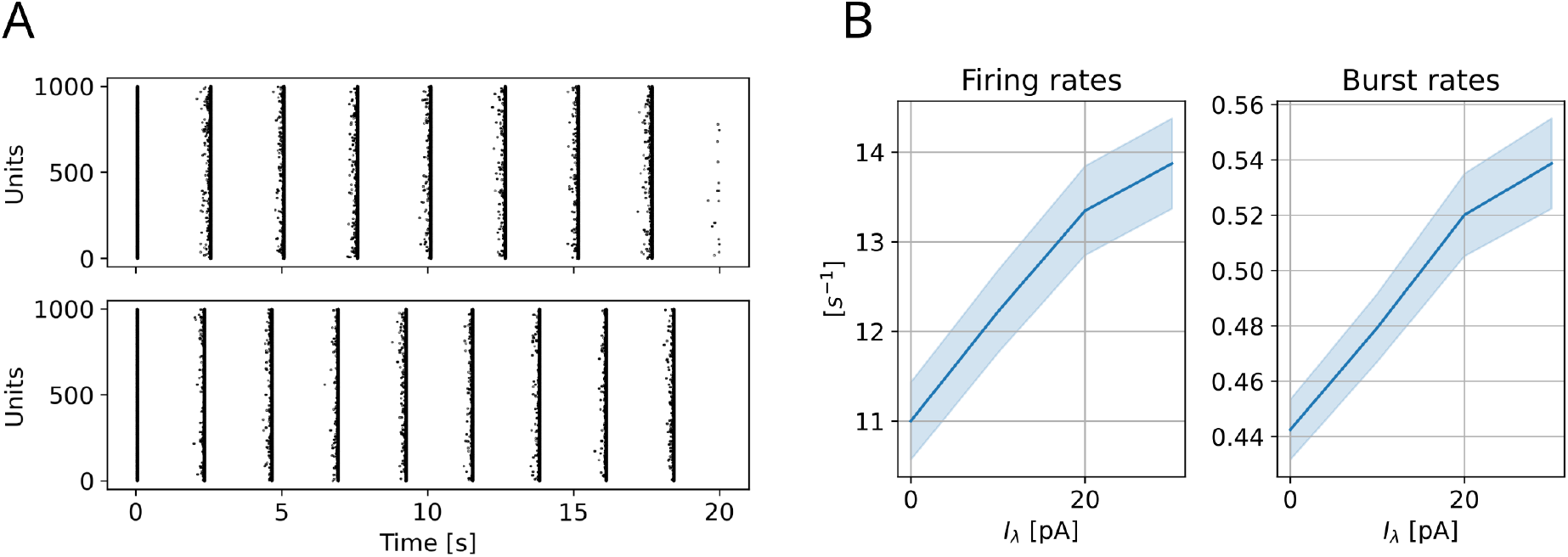
A: Simulated raster plot of the same network as in Fig. 2B for different values of *I*_*λ*_ in Eq. 14. Top: *I*_*λ*_ = 0pA. Bottom: *I*_*λ*_ = 10pA. B: Firing rate (left) and bursting rate (right) in the same network for different values of *I*_*λ*_. Solid line: Value averaged over all neurons and over 10 repetitions with random initial cell membranes. Shaded area: s.e.m.

Similar low-frequency oscillations had been previously observed in neuronal populations, especially during sleep, anesthesia, or in-vitro when using a mock cerebrospinal fluid (CSF) [48, 49]. Moreover, different computational models have been proposed to reproduce and explain them. In [48], the authors augment the classical Hodgkin-Huxley model with a slow sodium-dependent potassium current which mediates the transition between up and down states (respectively characterized by higher and lower levels of spontaneous activity). However, this model relies on an important population of inhibitory neurons to stabilize the strong recurrent excitation during the up state. In [50], the authors propose a simpler model (based on LIF neurons) to reproduce these slow-oscillations, but here again the recurrent excitation needs to be balanced by inhibitory neurons. This does not coincide with the very low proportion of inhibitory neurons in the observed population in [21].

**Table 2:** Parameter values used in Fig. 2 to Fig. 4. Values for *C*_*m*_, *g*_*L*_, *E*_*L*_, *V*_*T*_, Δ_*T*_, *a*, and *V*_*r*_ are the same as these used to simulate regular bursting in [53]. Values for *τ*_*w*_, *b*, and *I* were adjusted to fit the long inter-burst intervals observed in experimental data (Fig. 2A).

| Name | Value | Description |
| --- | --- | --- |
| $C_m$ | 200pF | Total capacitance |
| $g_L$ | 10nS | Total leak conductance |
| $E_L$ | -60mV | Effective rest potential |
| $\Delta_T$ | 2mV | Threshold slope factor |
| $V_T$ | -50mV | Effective threshold potential |
| $a$ | 2nS | Adaptation conductance |
| $\tau_w$ | 800mV | Adaptation time constant |
| $b$ | 30pA | Spike triggered adaptation |
| $V_r$ | -46mV | Reset potential |
| $I$ | 102pA | Injected current |
| $p$ | 0.02 | Connection probability |
| $w_e$ | 0.5mV | Post-synaptic potential amplitude |

However, other phenomena, such as firing rate adaptation [51] and short-term depression [52], have been proposed as ways to stabilize the coordinated activity of connected excitatory neurons without inhibitory feedback. In line with [53], we model the evolution of the voltage potential *V* in the presence of an injected current *I* using the adaptive Exponential Integrate-and-Fire model (AdEx), which can be seen as a simplification of Eq. 2 in that it neglects the individual ion-specific dynamics:

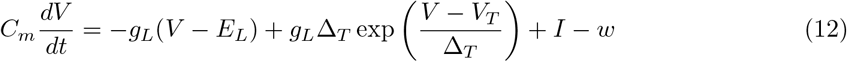

where *C*_*m*_ is the cell total membrane capacitance, *g*_*L*_ is the total leak conductance, *E*_*L*_ is the effective rest potential, Δ_*T*_ is the threshold slope factor, and *V*_*T*_ is the effective threshold potential. The last term *w* (which is not present in Eq. 2) tends to hyper-polarize the cell following a spike, hence reducing the probability of a subsequent spike and leading to spike-rate adaptation. It is increased by a quantity *b* following each spike, and otherwise decays with a time constant *τ*_*w*_:

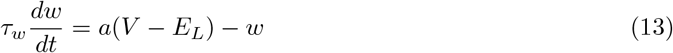

We verify that the stable self-sustained slow oscillations observed in neuronal cell lines (Fig. 2A) can be reproduced by a model consisting of a population of LIF neurons recurrently connected by excitatory synapses, whose activity is balanced by firing rate adaptation, without the need for an inhibitory population (Fig. 2B). Across all simulated neurons, the average firing rate was 10.85 *±* 1.39Hz and the average burst rate was 0.44 *±* 0.05Hz, which is in agreement with the experimental values reported in Figure 2D and E in [21].

### 2.4 The model reproduces features of network activity observed in altered gravity

Finally, after having identified the main features of the neuron networks used in altered gravity experiments (i.e. excitatory recurrent connections and population activity stabilized by firing rate adaptation), we verified that injecting an additional current onto this network (to represent the activity of MG channels) allows to reproduce the increased firing and burst rates observed during the dynamic phases of altered gravity experiments. This acceleration-induced current is represented by an extra term *I*_*λ*_ in the right-hand side of Eq. 12:

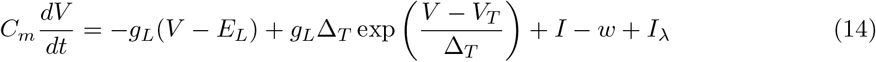

Fig. 3A shows the raster plot for the same network as in Fig. 2B, for *I*_*λ*_ = 0pA (top) and *I*_*λ*_ = 10pA (bottom). Fig. 3B shows the main firing rate (left) and burst rate (right) of the network (averaged over all neurons) as a function of *I*_*λ*_. Both the firing and burst rates increase with *I*_*λ*_, which is in line with experimental observations. More specifically, hypergravity experiments using the same hiPSC-derived neurons on the DLR human centrifuge in Cologne, Germany have reported a decreased neuronal activity compared to baseline recordings during exposure to 6g hypergravity (as detailed in section 2.2), but also an increased activity during both the ramp-up and ramp-down phases of the centrifuge (Fig. 2D and E in [21]). These ramp-up and ramp-down phases correspond to a dynamic modification of the perceived acceleration, during which MG channels are likely to be activated and to contribute to the depolarization of the cells.

In addition to its potential effect on ion channels dynamics (which was previously suggested in section 2.1), the gravity-induced modification of cellular membrane fluidity [23, 25, 36, 37] could also impact the activity of cells networks by modifying synaptic transmission. Indeed, both exocytosis and endocytosis, which are critical for synaptic transmission, are dependent on the fluidity of the cell membrane: a high tension will increase the exocytosis rate [56, 57] while also decreasing the endocytosis rate [58, 59]. These phenomena are still observed in the active zones of nerve terminals [60] and happen over fast time scale, making them potentially relevant for the rapid modification of neuronal activity following the onset of altered gravity. Since both endocytosis and exocytosis are critical for synaptic transmission, whether the decrease of the former and the increase of the latter (which are both induced by an increased membrane fluidity) will compensate each other, or whether one effect will dominate, remains an open question. However, it was also suggested that centrifugation hypergravity stress alters both neurotransmitter reuptake and exocytotic release of glutamate [61]: the effect of hypergravity (resp. microgravity) could thus be computationally represented as a decrease (resp. increase) of the connection weights in the simulated network. We added an offset 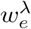 to the base post-synaptic potential amplitude *w*_*e*_. Interestingly, increasing the connection weights in the network increases its overall firing rate (Fig. 4A left), as observed in microgravity, but also reduces the bursting rate (Fig. 4A right): the inter-burst interval grows with 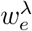 but so does the length of each burst (Fig. 4B), ultimately yielding an increased firing rate.

**Figure 4:**
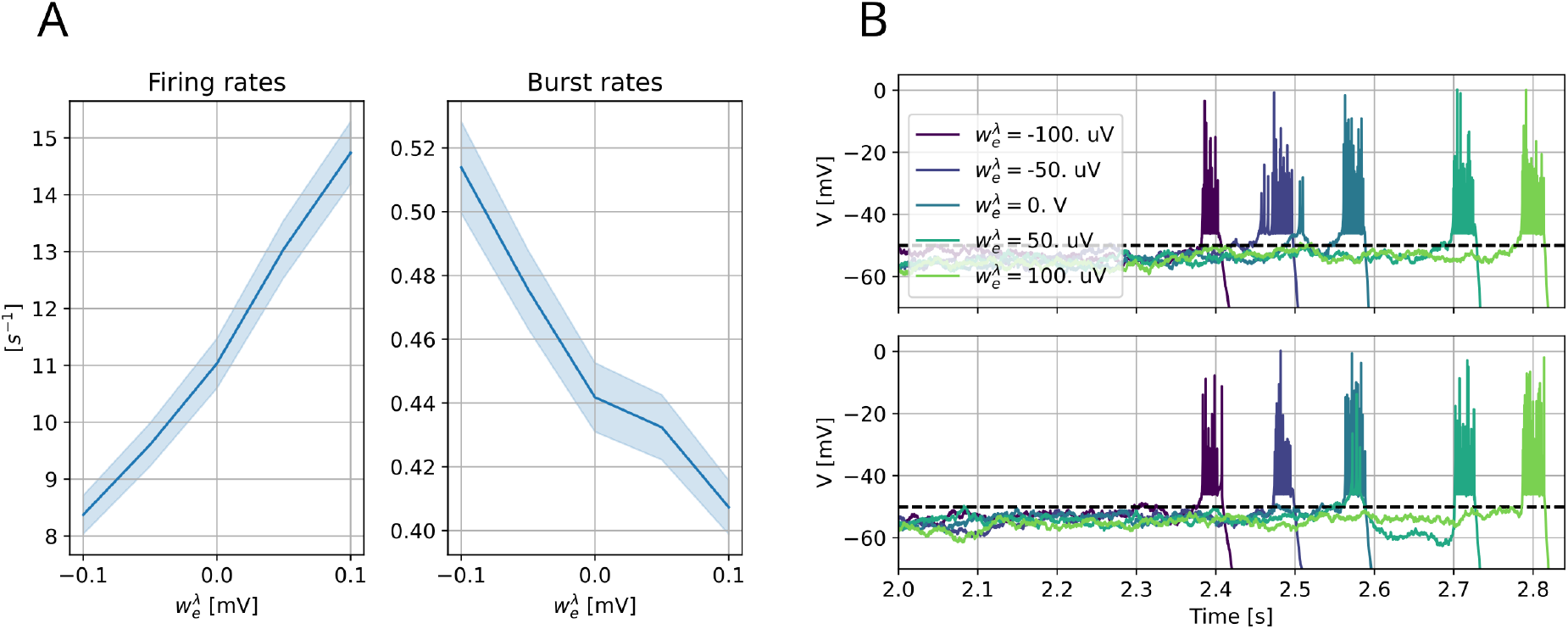
A: Firing rate (left) and bursting rate (right) in the same network as in Fig. 2B for different values of 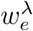 added to *w*_*e*_ in Table 2. Solid line: Value averaged over all neurons and over 10 repetitions with random initial cell membranes. Shaded area: s.e.m. B: Voltage traces for two example neurons and for different values of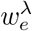, showing the increasing inter-burst interval with increasing values of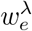.

**Figure 5:**
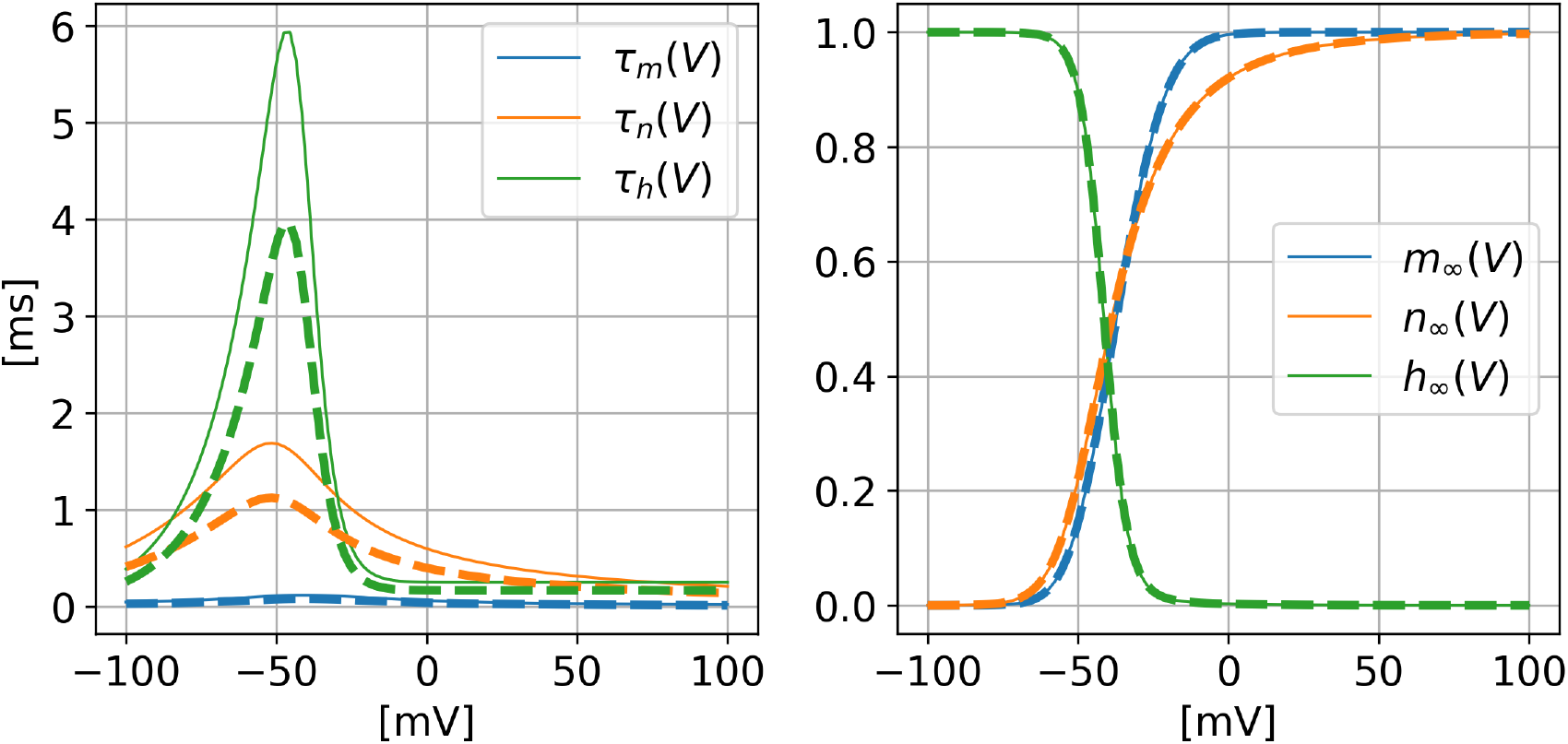
Effect of *λ* on the gating time constants (left) and steady-state levels (right) in the Hodgkin-Huxley model. Solid line: *λ* = 1. Dashed line: *λ* = 1.5.

**Figure 6:**
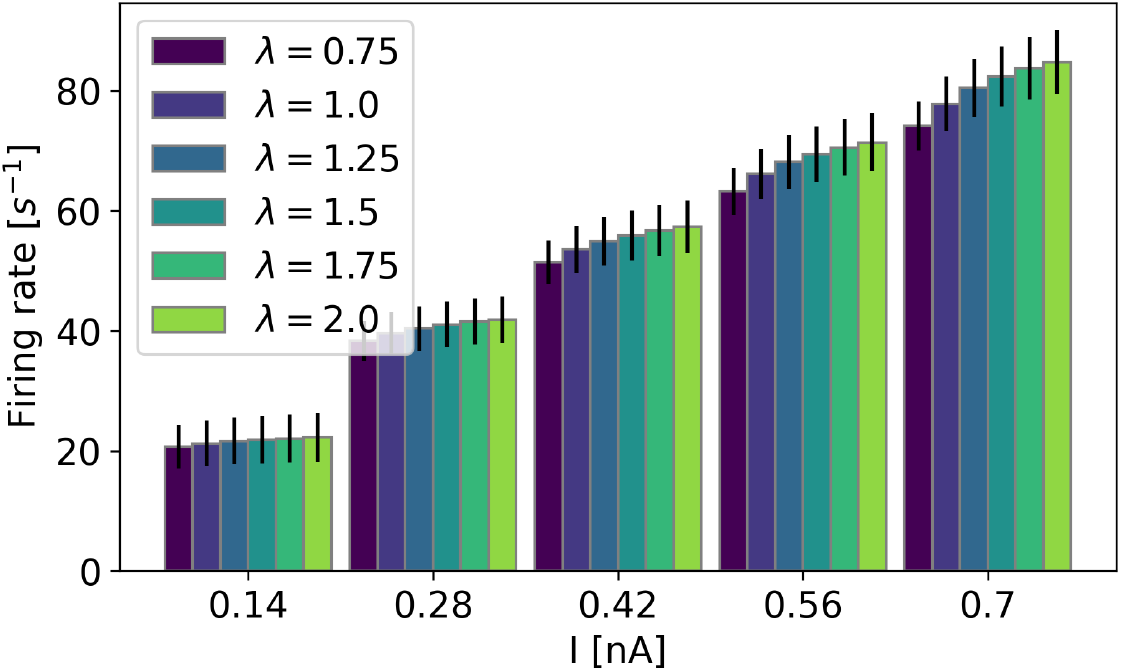
Robustness analysis: same results as in Fig. 1A, with results averaged over 100 independent repetitions. For each repetition, the value of each simulation parameter was randomly drawn from a uniform distribution with bounds *±*10% of the nominal values listed in Table 1. Vertical bars: s.e.m.

## 3 Discussion

Neuronal activity has been observed to be gravity-dependent. More specifically, MEA recordings have shown that the firing and burst rates of cultured stem cells increase during drop tower-induced microgravity and decrease during centrifuge-induced hypergravity, with important variability between neuron subtypes [21]. This is in line with calcium activity recordings of hippocampal neurons obtained during parabolic flights, which showed that more neurons are active during altered gravity than during the 1g baseline period, and that inter-spike intervals are shorter during microgravity phases compared to hypergravity phases [62].

However, literature on how to model this phenomenon is sparse. On the one hand, a qualitative model has been proposed, based on the influence of gravity on ion channels [23, 24]; but it neither accounts for correlated dynamics of ion channels nor for possible connectivity between neurons. On the other hand, classical computational models, such as the Hodgkin-Huxley model, or recurrent networks of in-silico neurons, do not have a gravity parameter that allows to represent the effect of altered gravity. Here, our approach is proposing computational implementations on how gravity impacts neuronal activity, which are coherent with both recorded neuronal activity under altered gravity and previous observations on how cell functions are gravity-dependent.

Many different in-silico parameters could be modified to yield a change in the firing and burst rates. However, a mechanism through which gravity alters neuronal activity should satisfy two points: First, it needs to be fast since modified neuronal activity upon altered gravity exposure happens on very short time scales [62]. Second, it should have been observed to happen during altered gravity experiments before. A first widely observed effect of gravity is on the fluidity of the cellular membrane [23, 25, 36, 37]. Based on it, a seminal explanatory model states that exposure to altered gravity modifies the physical properties of the cells’ membrane, ultimately altering their voltage dynamics [23, 24]. This is in line with previous theoretical studies which had already shown how, for the same stationary input current, firing rates can be modulated by the dynamical properties of the cell membrane to perform prospective decoding [63]. Since ion channel dynamics are gravity-dependent [33–35] and influenced by the properties of the cell membrane [38–44], we represent this effect by a linear scaling of the channels’ gating rate (Eqs. 9, 10, 11): we verify that it allows to reproduce altered firing and bursting rates, both in single neurons (Fig. 1A to C) and in connected networks (Fig. 1D).

Secondly, a previous study using centrifuge-induced hypergravity had reported an increase in neuronal activity both during the ramp up (i.e., the transition from 1g to hypergravity) and ramp down (i.e., the transition from hypergravity to 1g) phases. This effect was attributed to MG ion channels, which are present in neuronal cells [28] and can induce action potentials [29]. Stretching of cells during acceleration and deceleration phases might thus impact neuronal activity [21]. The effect of an additional excitatory current is easy to analyze for single cells, but may be not trivial in networks of connected neurons. We show that the main features of the cell lines used in altered gravity experiments (i.e., synchronized activity with low-frequency bursts) can be reproduced by a network of connected excitatory cells stabilized by firing rate adaptation (Fig. 2), and that its firing and bursting rates will increase upon addition of an excitatory current (Fig. 3).

Our overall contribution is several-fold. Based on previous studies, we identified mechanisms which may explain the fast neuronal responses to altered gravity (namely: the influence of membrane fluidity on ion channel dynamics and synaptic transmission, and the activation of depolarizing MG channels). We proposed simple computational implementations of these mechanisms, which yield results coherent with historical observations. Note that these identified phenomena are not mutually exclusive hypotheses, but potentially additive effects which future experiments can disentangle (see below). Our modeling assumptions suggest that these effects could be additive (see the increase in firing rates with both *I* and *λ* in Figs. 1A and 6). We address contradictory previous results on the effect of altered gravity on ion channels dynamics [23, 24, 33–35] and propose a unified explanation for the effect of altered gravity on membrane fluidity, ion channels dynamics, and neuronal firing rates. Finally, we fitted recordings with a network model including firing rate adaptation: this proposed computational model explains the activity of networks of neurons used in altered gravity experiments, and especially their coordinated, low-frequency burst patterns in the absence of inhibitory cells.

Our approach has evident limitations. Our single-neuron analysis is based on the Hodgkin-Huxley model, which is a middle point between the simplicity of LIF neurons (which do not model the dynamics of single channels) and the realism of biophysical neuron models (e.g. [64]). The Hodgkin-Huxley model is largely descriptive rather than mechanistic, and does not account for several mechanisms. Future work should consider extending our analysis to newer, more realistic neuron models (e.g. [48]).

Furthermore, the goal of our approach is not to provide a mechanism for the effect of gravity on neuronal activity. Rather, our approach is a bottom-up process: starting from prior observations on the effect of altered gravity on cellular functions, we proposed computational implementations which are coherent with the observed effect of altered gravity on neuronal firing rates. Nevertheless, our results could greatly benefit from ad-hoc intracellular recordings. Previous data on neuronal activity under altered gravity were obtained using either extracellular recordings [21] or calcium imaging [62]. A remote-controlled device for patch-clamping has been developed [65] and implemented on a drop tower and parabolic flights to study action potential propagation [66]. However, intracellular recordings under altered gravity remain rare due to the difficulty in carrying out such experiments and the sensitivity of the setup. Future experiments could be designed to isolate different effects, e.g. by silencing MG channels or modifying the experimental setup to follow potential changes in neuronal membrane fluidity. Patch clamp and sequencing experiments could help in further specifying the type and dynamics of the MG channels present, and to refine the assumptions used in our models.

The models we use in our simulations (and especially the Hodgkin-Huxley model) rely on a large number of free parameters (see Tables 1 and 2). However, these parameters can be easily constrained to small ranges of biologically plausible values. Seminal papers on conductance-based neurons [67] and firing rate adaptation [53] have proposed biologically plausible values for these parameters, which we based our simulations on. Some of these parameters (namely the adaptation parameters *τ*_*w*_ and *b*, and the connectivity parameters *I, p*, and *w*_*e*_ in Table 2) were adjusted to match the specificities of the studied neurons (and especially to reproduce the synchronized, low-frequency bursting activity observed in Fig. 2A and in [21]). We verified that these values are in line with these previously used in seminal studies on modeling neuron networks [47]. In Tables 1 and 2, the values for the capacitance *C*_*m*_ and leakage reversal potential *E*_*L*_ are in line with these reported for actual iNGN neuronal cells [68]. Future experiments could focus on estimating the exact value of the other parameters (i.e., the ion-specific parameters in Table 1, or the connectivity parameters in Table 2) for iNGN neurons using dedicated intracellular recordings. Overall, we argue that our results are based on realistic values of the simulation parameters, and that they are robust to reasonable variations in these values (Fig. 6).

The altered response observed in cell cultures usually immediately follows the onset of microgravity. However, in longer-term altered gravity experiments (i.e., involving sounding rockets), the neuronal activity recordings suggest a complex, dynamic response characterized by an initial increase followed by a transient decline and partial recovery over several minutes. The computational models outlined here only account for the former (i.e., the short-term modification of neuronal activity following exposure to microgravity, for which quantitative analyses and numerical verification were lacking). To account for the later (i.e., the relaxation of neuronal activity to baseline levels following prolonged exposure to altered gravity), many different phenomena and models have been proposed to explain response to activity perturbation, including individual neurons spike frequency adaptation [69–71] and synaptic homeostatic plasticity [72–74]. Further exploration into the dynamics of neuronal activity during prolonged exposure to microgravity will be essential to determine the persistence of changes in firing rates. Such long-term investigations could reveal implications for brain function, potentially leading to acute and chronic issues. More generally, whether results obtained at the cellular levels will translate at the behavioral and cognitive levels remains an open question. Recent studies performed during parabolic flights have reported a modification of the EEG signal, but not of the participants’ cognitive performances, under different micro- and hypergravity conditions [16, 75]: this hints at the existence of adaptation mechanisms, making cognition robust to altered gravity levels despite its effect on cultured cells.

Overall, our approach fills an important gap in the literature, by reviewing the mechanisms through which gravity levels could alter neuronal activity. In the absence of intracellular recordings obtained under various gravity levels, and following the principle of Occam’s razor, our computational implementations of these phenomena remain simple: linear scaling of the gating time constants, and an additional uniform excitatory current. Future work will focus on refining these modeling assumptions by obtaining intracellular recordings at different gravity levels.

## 4 Methods

### 4.1 Simulation of single neurons

In Fig. 1, the voltage of simulated neurons is governed by Eqs. 1, 2, 9, 10, and 11, with parameters values listed in Table 1. In Fig. 1D, we further implemented synaptic connections between simulated neurons. Following [67], the voltage of neuron *n* is given by

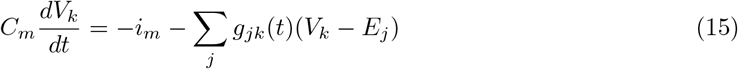

where *i*_*m*_ is given by Eq. 2 and *g*_*jk*_(*t*)(*V*_*k*_ − *E*_*j*_) represents the synaptic connection between neurons *k* and *j*. The value of the synaptic reversal potential *E*_*j*_ depends on whether the synapse is excitatory or inhibitory (see values in Table 1). The synaptic conductance *g*_*jk*_ decays exponentially with time constants *τ*_*e*_ (for excitatory synapses) and *τ*_*i*_ (for inhibitory synapses), and is incremented by a quantum *w*_*e*_ or *w*_*i*_ whenever an action potential is generated by neuron *j*.

### 4.2 Simulation of networks of neurons

In Figs. 2 to 4, the evolution of the voltage of each individual neuron is governed by Eqs. 12 to 14, with parameters values listed in Table 2. Furthermore, simple voltage-jump synapses are implemented as follows: a synaptic connection from neuron *j* to *k* is randomly created with probability *p*. If a synaptic connection is established, the voltage of neuron *k* is immediately increased by a quantum *w*_*e*_ whenever neuron *j* generates an action potential.

### 4.3 Experimental data

This study mostly refers to previously analyzed and published data. Only in Fig. 2A are original data from a recent rocket flight presented for illustration purposes.

Methods and procedures that were employed to collect the data were previously described in [21]. In brief, we used human induced pluripotent stem cell (hiPSC)-derived neurons (iNGN cells) to conduct the experiments. Upon induction via the TetOn system, Neurogenin-1 and Neurogenin-2 were expressed and cells differentiated into post-mitotic, bipolar neurons [76]. iNGN cells were cultured and reseeded on MEAs following a previously published protocol [77]. Thawed iNGN cells were cultured on Matrigel-coated plates in mTeSR™1 medium (STEMCELL Technologies, Germany). After at least two passages, cells were induced with 0.5 *µ*g/ml doxycycline for three days. At 2 dpi, 5 *µ*M Ara-C was added to eliminate undifferentiated cells. Concurrently, MEAs were prepared with poly-D-lysine and laminin coatings. Induced cells were reseeded on MEA chips at 3 dpi. Cells were dissociated, centrifuged, and resuspended in BrainPhys™ medium (STEMCELL Technologies, Germany), and 100,000 cells were seeded per MEA. BrainPhys™ medium was supplemented with neurotrophic factors and ascorbic acid. Astrocyte-conditioned media was added for long-term culture. Media was partially refreshed weekly with a 1:1 mix of fresh and astrocyte-conditioned BrainPhys™ medium.

The experimental setup included a MEA system housed in a pressure chamber to maintain physiological conditions. The MEA’s integrated heaters kept cells at 37°C, while the pressure-tight chamber ensured stable pressure and CO2 levels. The system’s compact design is compatible with various altered gravity research platforms (DLR SAHC, ZARM drop tower, parabolic flights, and sounding rockets). A rail system allows for easy removal of the experiment module, facilitating time-sensitive operations like those on sounding rockets.

After culturing MEA samples for several weeks, they were brought to respective experiment facilities using transport incubators (CellTrans 2018 and 4016, Labotect, Germany; 37°C, 5% CO2).

Launching rocket experiments were carried out at the ESA ESRANGE space center in Kiruna, Sweden. iNGN MEA cultures were transported to the experiment site at least 10 days before launch. During the hot countdown and right before late access, MEA chips were integrated into the BIODECODER module and functionality of the system was checked. The entire module was kept under a constant temperature of 37°C within a custom-designed transport incubator (DLR). Upon late access, which is a time window in between 2.5 to 1 h before launch, the BIODECODER module containing the samples was loaded into the dedicated slot within the rocket. Recording was started 10 min prior to launch and data was recorded continuously throughout the entire flight, including post-baseline recording back on ground. Battery operation ensured constant heating of the samples after launch and landing. The module was retrieved by a helicopter and brought back to the launch site, where additional post-experiment recordings and microscopy were executed.

## Acknowledgments

We thank Martin Müller, Nicolas Vitale, Jakob Jordan, Aitor Morales- Gregorio, Igor Delvendahl, Roberto Martins de Freitas, Denis-Gabriel Caprace, Mohammad Iran- manesh, Mehdi Scoubeau, Simon Benjamin Brandt, Timo Gierlich, Ben von Hünerbein, and Jérémy Rabineau for the fruitful discussions. This study was funded by the German Aerospace Center. JS acknowledges support by the Joachim Herz Foundation. The funders played no role in study design, data collection, analysis and interpretation of data, or the writing of this manuscript.

## Supplementary material

**Figure 7:**
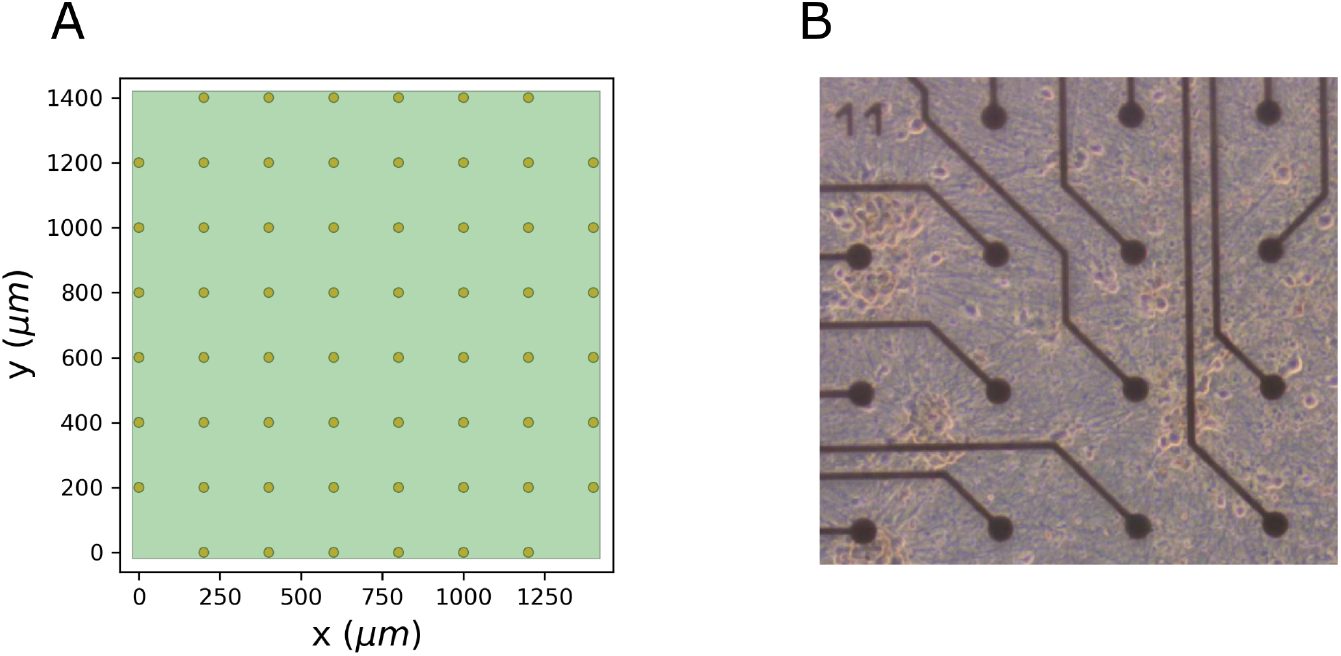
A: Layout of the recording electrodes. Population activity was recorded using Multi-Electrode Array (MEA) chips featuring 60 individual electrodes (60MEA200/30iR-ITO chips, MultiChannel Systems, Germany). The 60 electrodes have a diameter of 30*µ*m and are separated by 200*µ*m. The extracellular voltage recorded by each electrode is sampled at 25 kHz. Each neuronal population was recorded using 2 sets of MEA chips simultaneously. B: Zoomed phase contrast microscopy image of an exemplary MEA chip showing different neurons. Reproduced from [21].

## Footnotes

1 Although the lipid composition is more important than bulk membrane fluidity for ion channel function [42].

## References

[1] P. E. di Prampero and M. V. Narici, “Muscles in microgravity: From fibres to human motion,” Journal of Biomechanics, vol. 36, no. 3, pp. 403–412, 2003.

[2] P. Droppert, “A review of muscle atrophy in microgravity and during prolonged bed rest.,” Journal of the British Interplanetary Society, vol. 46, no. 3, pp. 83–86, 1993.

[3] D. Grimm et al., “The impact of microgravity on bone in humans,” Bone, vol. 87, pp. 44–56, 2016.

[4] A. R. Hargens and S. Richardson, “Cardiovascular adaptations, fluid shifts, and countermeasures related to space flight,” Respiratory Physiology & Neurobiology, vol. 169, S30–S33, 2009.

[5] J. S. Lawley et al., “Effect of gravity and microgravity on intracranial pressure,” The Journal of Physiology, vol. 595, no. 6, pp. 2115–2127, 2017.

[6] A. G. Lee, T. H. Mader, C. R. Gibson, T. J. Brunstetter, and W. J. Tarver, “Space flight-associated neuro-ocular syndrome (sans),” Eye, vol. 32, no. 7, pp. 1164–1167, 2018.

[7] A. G. Lee et al., “Spaceflight associated neuro-ocular syndrome (sans) and the neuro-ophthalmologic effects of microgravity: A review and an update,” npj Microgravity, vol. 6, no. 1, pp. 1–10, 2020.

[8] D. Roberts et al., “Prolonged microgravity affects human brain structure and function,” American Journal of Neuroradiology, vol. 40, no. 11, pp. 1878–1885, 2019.

[9] N. K. Popova, A. V. Kulikov, and V. S. Naumenko, “Spaceflight and brain plasticity: Spaceflight effects on regional expression of neurotransmitter systems and neurotrophic factors encoding genes,” Neuroscience & Biobehavioral Reviews, vol. 119, pp. 396–405, 2020.

[10] R. Herranz et al., “Ground-based facilities for simulation of microgravity: Organism-specific recommendations for their use, and recommended terminology,” Astrobiology, vol. 13, no. 1, pp. 1–17, 2013.

[11] J. Hauslage et al., “Cytosolic calcium concentration changes in neuronal cells under clinorotation and in parabolic flight missions,” Microgravity Science and Technology, vol. 28, pp. 633– 638, 2016.

[12] N. Callens, J. Ventura-Traveset, T.-L. de Lophem, C. Lopez de Echazarreta, V. Pletser, and J. J. van Loon, “Esa parabolic flights, drop tower and centrifuge opportunities for university students,” Microgravity Science and Technology, vol. 23, pp. 181–189, 2011.

[13] C. Liemersdorf, Y. Lichterfeld, R. Hemmersbach, and J. Hauslage, “The mapheus module cellfix for studying the influence of altered gravity on the physiology of single cells,” Review of Scientific Instruments, vol. 91, no. 1, 2020.

[14] J. Hauslage et al., “Arabidomics—a new experimental platform for molecular analyses of plants in drop towers, on parabolic flights, and sounding rockets,” Review of Scientific Instruments, vol. 91, no. 3, 2020.

[15] V. Pletser, “Short duration microgravity experiments in physical and life sciences during parabolic flights: The first 30 esa campaigns,” Acta Astronautica, vol. 55, no. 10, pp. 829– 854, 2004.

[16] C. Badali, P. Wollseiffen, and S. Schneider, “Shades of gravity–effects of planetary gravity levels on electrocortical activity and neurocognitive performance,” Brain Structure and Function, pp. 1–13, 2024.

[17] A. Acharya et al., “Parabolic, flight-induced, acute hypergravity and microgravity effects on the beating rate of human cardiomyocytes,” Cells, vol. 8, no. 4, p. 352, 2019.

[18] H. Selig, A. Gierse, and G. König, “Parabolic flight with light aircraft,” in 67th International Astronautical Congress (IAC). International Astronautical Federation (IAF), Guadalajara.(IAC16-A2-5-6-x34402), 2016.

[19] D.-G. Caprace, C. Gontier, M. Iranmanesh, M. Scoubeau, and V. Pletser, “Experimental characterization of weightlessness during glider parabolic flights,” Microgravity Science and Technology, vol. 32, no. 6, pp. 1121–1132, 2020.

[20] Y. Lichterfeld et al., “Hypergravity attenuates reactivity in primary murine astrocytes,” Biomedicines, vol. 10, no. 8, p. 1966, 2022.

[21] J. Striebel et al., “Human neural network activity reacts to gravity changes in vitro,” Frontiers in Neuroscience, vol. 17, p. 1 085 282, 2023.

[22] V. Pletser, N. Frischauf, R. Laufer, and D. Cohen, “Parabolic flights with gliders as an innovative low-cost platform for microgravity and hypergravity research,” in 68th International Astronautical Congress (IAC). International Astronautical Federation (IAF), Adelaide,.(IAC17-A2-5-6-x36752), 2017, pp. 1–5.

[23] F. Kohn, J. Hauslage, and W. Hanke, “Membrane fluidity changes, a basic mechanism of interaction of gravity with cells?” Microgravity Science and Technology, vol. 29, pp. 337–342, 2017.

[24] F. P. Kohn and R. Ritzmann, “Gravity and neuronal adaptation, in vitro and in vivo—from neuronal cells up to neuromuscular responses: A first model,” European Biophysics Journal, vol. 47, pp. 97–107, 2018.

[25] F. P. Kohn and J. Hauslage, “The gravity dependence of pharmacodynamics: The integration of lidocaine into membranes in microgravity,” npj Microgravity, vol. 5, no. 1, p. 5, 2019.

[26] D.-P. Häder, M. Braun, D. Grimm, and R. Hemmersbach, “Gravireceptors in eukaryotes—a comparison of case studies on the cellular level,” npj Microgravity, vol. 3, no. 1, p. 13, 2017.

[27] S. Kirischuk, “Mechanosensitive channels in neuronal and astroglial cells in the nervous system,” Mechanosensitivity of the Nervous System: Forewords by Nektarios Tavernarakis and Pontus Persson, pp. 3–22, 2009.

[28] F. Falleroni et al., “Mechanotransduction in hippocampal neurons operates under localized low piconewton forces,” Iscience, vol. 25, no. 2, 2022.

[29] Y. A. Nikolaev, P. J. Dosen, D. R. Laver, D. F. Van Helden, and O. P. Hamill, “Single mechanically-gated cation channel currents can trigger action potentials in neocortical and hippocampal pyramidal neurons,” Brain Research, vol. 1608, pp. 1–13, 2015.

[30] R. Habibey, J. Striebel, F. Schmieder, J. Czarske, and V. Busskamp, “Long-term morphological and functional dynamics of human stem cell-derived neuronal networks on high-density microelectrode arrays,” Frontiers in Neuroscience, vol. 16, p. 951 964, 2022.

[31] A. L. Hodgkin and A. F. Huxley, “A quantitative description of membrane current and its application to conduction and excitation in nerve,” The Journal of Physiology, vol. 117, no. 4, p. 500, 1952.

[32] P. Dayan and L. F. Abbott, Theoretical neuroscience: computational and mathematical modeling of neural systems. MIT Press, 2005.

[33] N. Klinke, M. Goldermann, and W. Hanke, “The properties of alamethicin incorporated into planar lipid bilayers under the influence of microgravity,” Acta Astronautica, vol. 47, no. 10, pp. 771–773, 2000.

[34] W. Hanke, “Studies of the interaction of gravity with biological membranes using alamethicin doped planar lipid bilayers as a model system,” Advances in Space Research, vol. 17, no. 6-7, pp. 143–150, 1996.

[35] M. Goldermann and W. Hanke, “Ion channel are sensitive to gravity changes,” Microgravity science and technology, vol. 13, no. 1, pp. 35–38, 2001.

[36] R. Asuwin Prabu, A. Manohar, S. Narendran, A. Kabir, and S. Sudhakar, “Effect of simulated microgravity on artificial single cell membrane mechanics,” Microgravity Science and Technology, vol. 36, no. 4, p. 47, 2024.

[37] D. Klymchuk, V. Baranenko, T. Vorobyova, I. Kurylenko, O. Chyzhykova, and V. Dubovoy, “Properties of plasma membrane from pea root seedlings under altered gravity,” in 35th COSPAR Scientific Assembly, vol. 35, 2004, p. 1356.

[38] C. Dart, “Symposium review: Lipid microdomains and the regulation of ion channel function,” The Journal of physiology, vol. 588, no. 17, pp. 3169–3178, 2010.

[39] A. L. Duncan, W. Song, and M. S. Sansom, “Lipid-dependent regulation of ion channels and g protein–coupled receptors: Insights from structures and simulations,” Annual review of pharmacology and toxicology, vol. 60, no. 1, pp. 31–50, 2020.

[40] M. L. Roberts-Crowley, T. Mitra-Ganguli, L. Liu, and A. R. Rittenhouse, “Regulation of voltage-gated ca2+ channels by lipids,” Cell calcium, vol. 45, no. 6, pp. 589–601, 2009.

[41] P. A. Schmidpeter et al., “Membrane phospholipids control gating of the mechanosensitive potassium leak channel trek1,” Nature communications, vol. 14, no. 1, p. 1077, 2023.

[42] C. Sunshine and M. G. McNamee, “Lipid modulation of nicotinic acetylcholine receptor function: The role of membrane lipid composition and fluidity,” Biochimica et Biophysica Acta (BBA)-Biomembranes, vol. 1191, no. 1, pp. 59–64, 1994.

[43] C. Sunshine and M. G. McNamee, “Lipid modulation of nicotinic acetylcholine receptor function: The role of neutral and negatively charged lipids,” Biochimica Et Biophysica Acta (BBA)-Biomembranes, vol. 1108, no. 2, pp. 240–246, 1992.

[44] L. Zanello, E. Aztiria, S. Antollini, and F. Barrantes, “Nicotinic acetylcholine receptor channels are influenced by the physical state of their membrane environment,” Biophysical journal, vol. 70, no. 5, pp. 2155–2164, 1996.

[45] M. Stimberg, R. Brette, and D. F. Goodman, “Brian 2, an intuitive and efficient neural simulator,” eLife, vol. 8, F. K. Skinner, Ed., e47314, Aug. 2019, issn: 2050-084X. doi: 10.7554/eLife.47314.

[46] C. Lu et al., “Overexpression of neurog2 and neurog1 in human embryonic stem cells produces a network of excitatory and inhibitory neurons,” The FASEB Journal, vol. 33, no. 4, p. 5287, 2019.

[47] B. Kriener, H. Enger, T. Tetzlaff, H. E. Plesser, M.-O. Gewaltig, and G. T. Einevoll, “Dynamics of self-sustained asynchronous-irregular activity in random networks of spiking neurons with strong synapses,” Frontiers in Computational Neuroscience, vol. 8, p. 136, 2014.

[48] A. Compte, M. V. Sanchez-Vives, D. A. McCormick, and X.-J. Wang, “Cellular and network mechanisms of slow oscillatory activity (¡ 1 hz) and wave propagations in a cortical network model,” Journal of Neurophysiology, vol. 89, no. 5, pp. 2707–2725, 2003.

[49] C. Capone, E. Pastorelli, B. Golosio, and P. S. Paolucci, “Sleep-like slow oscillations improve visual classification through synaptic homeostasis and memory association in a thalamo-cortical model,” Scientific Reports, vol. 9, no. 1, p. 8990, 2019.

[50] M. Torao-Angosto, A. Manasanch, M. Mattia, and M. V. Sanchez-Vives, “Up and down states during slow oscillations in slow-wave sleep and different levels of anesthesia,” Frontiers in Systems Neuroscience, vol. 15, p. 609 645, 2021.

[51] B. Gutkin and F. Zeldenrust, “Spike frequency adaptation,” Scholarpedia, vol. 9, no. 2, p. 30 643, 2014, revision #143322. doi: 10.4249/scholarpedia.30643.

[52] C. v. Vreeswijk and D. Hansel, “Patterns of synchrony in neural networks with spike adaptation,” Neural Computation, vol. 13, no. 5, pp. 959–992, 2001.

[53] R. Naud, N. Marcille, C. Clopath, and W. Gerstner, “Firing patterns in the adaptive exponential integrate-and-fire model,” Biological Cybernetics, vol. 99, pp. 335–347, 2008.

[54] A. P. Buccino et al., “Spikeinterface, a unified framework for spike sorting,” eLife, vol. 9, e61834, 2020.

[55] J. E. Chung et al., “A fully automated approach to spike sorting,” Neuron, vol. 95, no. 6, pp. 1381–1394, 2017.

[56] N. C. Gauthier, M. A. Fardin, P. Roca-Cusachs, and M. P. Sheetz, “Temporary increase in plasma membrane tension coordinates the activation of exocytosis and contraction during cell spreading,” Proceedings of the National Academy of Sciences, vol. 108, no. 35, pp. 14 467– 14 472, 2011.

[57] T. A. Masters, B. Pontes, V. Viasnoff, Y. Li, and N. C. Gauthier, “Plasma membrane tension orchestrates membrane trafficking, cytoskeletal remodeling, and biochemical signaling during phagocytosis,” Proceedings of the National Academy of Sciences, vol. 110, no. 29, pp. 11 875– 11 880, 2013.

[58] J. Dai, H. P. Ting-Beall, and M. P. Sheetz, “The secretion-coupled endocytosis correlates with membrane tension changes in rbl 2h3 cells,” The Journal of general physiology, vol. 110, no. 1, pp. 1–10, 1997.

[59] D. Raucher and M. P. Sheetz, “Membrane expansion increases endocytosis rate during mitosis,” The Journal of cell biology, vol. 144, no. 3, pp. 497–506, 1999.

[60] X.-S. Wu et al., “Membrane tension inhibits rapid and slow endocytosis in secretory cells,” Biophysical journal, vol. 113, no. 11, pp. 2406–2414, 2017.

[61] T. Borisova, N. Krisanova, and N. Himmelreich, “Exposure of animals to artificial gravity conditions leads to the alteration of the glutamate release from rat cerebral hemispheres nerve terminals,” Advances in Space Research, vol. 33, no. 8, pp. 1362–1367, 2004.

[62] P.-E. Lecoq, C. Dupuis, X. Mousset, X. Benoit-Gonnin, J.-M. Peyrin, and J.-L. Aider, “Influence of microgravity on spontaneous calcium activity of primary hippocampal neurons grown in microfluidic chips,” npj Microgravity, vol. 10, no. 1, p. 15, 2024.

[63] S. Brandt, M. A. Petrovici, W. Senn, K. A. Wilmes, and F. Benitez, Prospective and retrospective coding in cortical neurons, 2024. 2405.14810 [q-bio.NC].

[64] M. Deistler et al., “Differentiable simulation enables large-scale training of detailed biophysical models of neural dynamics,” bioRxiv, pp. 2024–08, 2024.

[65] K. Meissner and W. Hanke, “Patch-clamp experiments under micro-gravity,” in Life in Space for Life on Earth, vol. 501, 2002, pp. 399–400.

[66] K. Meissner and W. Hanke, “Action potential properties are gravity dependent,” Microgravity Science and Technology, vol. 17, pp. 38–43, 2005.

[67] R. Brette et al., “Simulation of networks of spiking neurons: A review of tools and strategies,” Journal of Computational Neuroscience, vol. 23, pp. 349–398, 2007.

[68] R. S. Lam, F. M. Töpfer, P. G. Wood, V. Busskamp, and E. Bamberg, “Functional maturation of human stem cell-derived neurons in long-term cultures,” PloS One, vol. 12, no. 1, e0169506, 2017.

[69] J. Benda and A. V. Herz, “A universal model for spike-frequency adaptation,” Neural Computation, vol. 15, no. 11, pp. 2523–2564, 2003.

[70] G. La Camera, A. Rauch, D. Thurbon, H.-R. Luscher, W. Senn, and S. Fusi, “Multiple time scales of temporal response in pyramidal and fast spiking cortical neurons,” Journal of Neurophysiology, vol. 96, no. 6, pp. 3448–3464, 2006.

[71] G. Fuhrmann, H. Markram, and M. Tsodyks, “Spike frequency adaptation and neocortical rhythms,” Journal of Neurophysiology, vol. 88, no. 2, pp. 761–770, 2002.

[72] I. Delvendahl and M. Müller, “Homeostatic plasticity—a presynaptic perspective,” Current Opinion in Neurobiology, vol. 54, pp. 155–162, 2019.

[73] C. A. Frank, T. D. James, and M. Müller, “Homeostatic control of drosophila neuromuscular junction function,” Synapse, vol. 74, no. 1, e22133, 2020.

[74] G. W. Davis and M. Müller, “Homeostatic control of presynaptic neurotransmitter release,” Annual Review of Physiology, vol. 77, pp. 251–270, 2015.

[75] C. Badali, P. Wollseiffen, and S. Schneider, “Under pressure—the influence of hypergravity on electrocortical activity and neurocognitive performance,” Experimental Brain Research, vol. 241, no. 9, pp. 2249–2259, 2023.

[76] V. Busskamp et al., “Rapid neurogenesis through transcriptional activation in human stem cells,” Molecular systems biology, vol. 10, no. 11, p. 760, 2014.

[77] F. Schmieder, R. Habibey, J. Striebel, L. Büttner, J. Czarske, and V. Busskamp, “Tracking connectivity maps in human stem cell–derived neuronal networks by holographic optogenetics,” Life Science Alliance, vol. 5, no. 7, 2022.

